# The genome regulatory landscape of Atlantic salmon liver through smoltification

**DOI:** 10.1101/2023.08.16.553484

**Authors:** Thomas N. Harvey, Gareth B. Gillard, Line L. Røsæg, Fabian Grammes, Øystein Monsen, Jon Olav Vik, Torgeir R. Hvidsten, Simen R. Sandve

## Abstract

The anadromous Atlantic salmon undergo a preparatory physiological transformation before seawater entry, referred to as smoltification. Key molecular developmental processes involved in this life stage transition, such as remodeling of gill functions, are known to be synchronized and modulated by environmental cues like photoperiod. However, little is known about the photoperiod influence and genome regulatory processes driving other canonical aspects of smoltification such as the large-scale changes in lipid metabolism and energy homeostasis in the developing smolt liver.

Here we generate transcriptome, DNA methylation, and chromatin accessibility data from salmon livers across smoltification under different photoperiod regimes. We find a systematic reduction of expression levels of genes with a metabolic function, such as lipid metabolism, and increased expression of energy related genes such as oxidative phosphorylation, during smolt development in freshwater. However, in contrast to similar studies of the gill, smolt liver gene expression prior to seawater transfer was not impacted by photoperiodic history. Integrated analyses of gene expression and transcription factor (TF) binding signatures highlight likely important TF dynamics underlying smolt gene regulatory changes. We infer that ZNF682, KLFs, and NFY TFs are important in driving a liver metabolic shift from synthesis to break down of organic compounds in freshwater. Moreover, the increased expression of ribosomal associated genes after smolts were transferred to seawater was associated with increased occupancy of NFIX and JUN/FOS TFs proximal to transcription start sites, which could be the molecular consequence of rising levels of circulating growth hormones after seawater transition. We also identified differential methylation patterns across the genome, but associated genes were not functionally enriched or correlated to observed gene expression changes across smolt development. This contrasts with changes in TF binding which were highly correlated to gene expression, underscoring the relative importance of chromatin accessibility and transcription factor regulation in smoltification.

**Author summary:** Atlantic salmon migrate between freshwater and seawater as they mature and grow. To survive the transition between these distinct environments, salmon transform their behavior, morphology, and physiology through the process of smoltification. One important adaptation to life at sea is remodeling of metabolism in the liver. It is unknown, however, whether this is a preadaptation that occurs before migration, what degree this is influenced by day length like other aspects of smoltification, and how gene regulatory programs shift to accomplish this transformation. We addressed these questions through a time course experiment where salmon were exposed to short and long day lengths, smoltified, and transferred to seawater. We sampled the livers and measured changes in gene expression, DNA methylation, chromatin accessibility, and transcription factor binding. We found metabolic remodeling occurred in freshwater before exposure to seawater and that day length did not have any long-term effects in liver. Transcription factor binding dynamics were closely linked to gene expression changes, and we describe transcription factors with key roles in smoltification. In stark contrast, we found no links between gene expression changes and DNA methylation patterns. This work deepens our understanding of the regulatory gear shifts associated with metabolic remodeling during smoltification.

## Introduction

Atlantic salmon are an anadromous species. They begin life in freshwater riverine habitats, then migrate to sea to grow and mature before returning to freshwater to spawn. The seawater migration is preceded by a “preparatory” process that influences a range of behavioral, morphological and physiological traits, referred to as smoltification [1]. This includes changes in pigmentation and growth [2], ion regulation [3, 4], the immune system [5], and various functions of the metabolism [6, 7].

The timing of smoltification is regulated by the physiological status of the fish [8], as well as external environmental signals such as temperature and day length [2, 9, 10]. Salmon smoltify in the spring, and the transition from short to long days is believed to drive changes in hormonal regulation and initiate smoltification. In line with this model, we recently demonstrated that exposure to a short photoperiod (i.e. a simulated winter photoperiod) induce transcription of a subset of photoperiod-history sensitive genes [3], dampens acute transcriptomic responses to increased salinity, and results in enhanced seawater growth [11]. These findings support a model of smolt development regulation, where photoperiodic-history drives genome regulatory remodeling underlying key smoltification associated phenotypes.

Although gill physiology has received most attention in the smoltification literature, other organs such as the liver also undergo large changes in function upon smoltification and seawater migration, with large implications for key metabolic traits. It has been shown that lipid composition in Atlantic salmon reared on different diets converges after smoltification [12, 13]. This is likely a consequence of smoltification associated increase in lipolytic rates and decreased lipid biosynthesis [6, 7]. In a recent study we demonstrated large changes in lipid metabolism gene regulation across the fresh-saltwater transition following smoltification [14]. Unfortunately, in this study smoltification and seawater transfer were confounded (i.e. smolts in freshwater were not sampled), hence it remains unclear if photoperiodic history is involved in shaping the molecular phenotype of the smolt liver as we observe in gills.

In this study, we conducted a smoltification trial to test if the photoperiodic history is a major factor impacting the genome regulatory landscape of Atlantic salmon liver. To do this we generated transcriptome, chromatin accessibility, and DNA methylation data across the smolt development and seawater transfer to characterize the transcriptomic changes in smolts reared with a short winter-like photoperiod (8:16) compared to smolts reared on constant light (24:0). We test if photoperiodic history affects the smolt liver phenotype at the level of gene expression and use chromatin accessibility data to identify putative regulatory pathways and transcription factors involved in life-stage associated changes in liver function from the juvenile stage in the freshwater environment to an adult fish in seawater.

## Results

### Gene expression changes support decreased lipid metabolism and increased protein metabolism and energy production during smoltification

A main goal was determining the effects of smoltification on metabolism and whether there was an effect of exposure to a short photoperiod (i.e. a winter) on the gene regulation in the liver. To accomplish this, we reared three groups of salmon for 46 weeks on commercial diets, from parr, through smoltification, and 6 weeks following transfer to seawater (Fig 1). The experimental group was given an artificial winter-like short photoperiod (8 hours light, 16 hours dark) for 8 weeks before they were returned to constant light, while the control group was reared under constant light throughout the experiment. Finally, the freshwater control group contained fish from the experimental group that was not transferred to sea. Following smoltification, fish transferred to seawater grew more slowly than fish that remained in freshwater (Fig 1, Table S1). There was no mortality throughout the freshwater portion of the trial, but some mortality (8x fish) in one tank due to improper oxygenation after seawater transfer.

**Fig 1:**
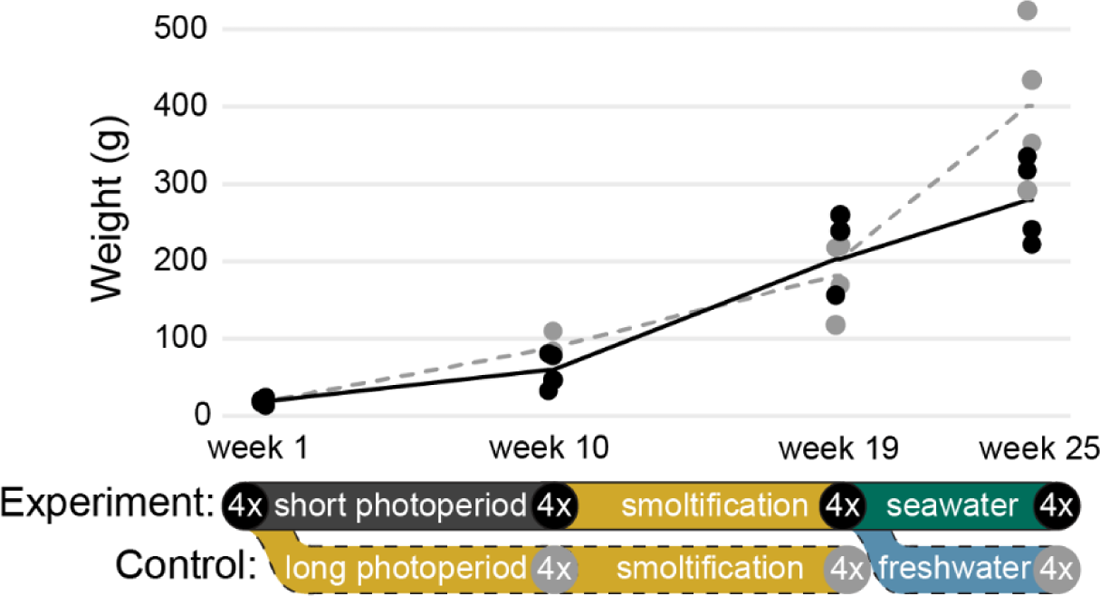
Salmon growth over time. Schematic of the experimental design and weight of salmon over time. Fish were reared for 21 weeks after first feeding in constant light conditions prior to week 1 sampling. The experimental group (black, solid line) was exposed to a short photoperiod before switched back to constant light and sampled at week 10. After a smoltification period, fish were sampled at week 19, then transferred to seawater conditions and sampled lastly at week 25. A photoperiod control group (grey, dashed line) received constant light throughout the experiment, and a freshwater control group branched off from the experimental group by remaining in freshwater. Four fish were sampled at each timepoint.

To characterize global transcriptome changes through key life stages, under a semi-natural developmental trajectory, we sampled liver tissue from fish at each sampling point for RNA sequencing. We first tested for changes in gene expression in fish experiencing artificial winter and transfer to seawater (experimental group) using an ANOVA-like test. This yielded 3,845 differentially expressed genes (DEGs, FDR <0.05) which were assigned to seven co-expression clusters using hierarchical clustering (Fig 2A, Table S2). These clusters reflected major patterns of gene regulatory changes (Fig 2B); peak expression levels in smolts (clusters 2 and 3), peak expression following the short photoperiod (clusters 4 and 5), decreased expression after short photoperiod and in smolts relative to all other time points (cluster 6), steady decrease in expression from parr throughout the experiment (cluster 7), and strong increase in expression in seawater (cluster 1).

**Fig 2:**
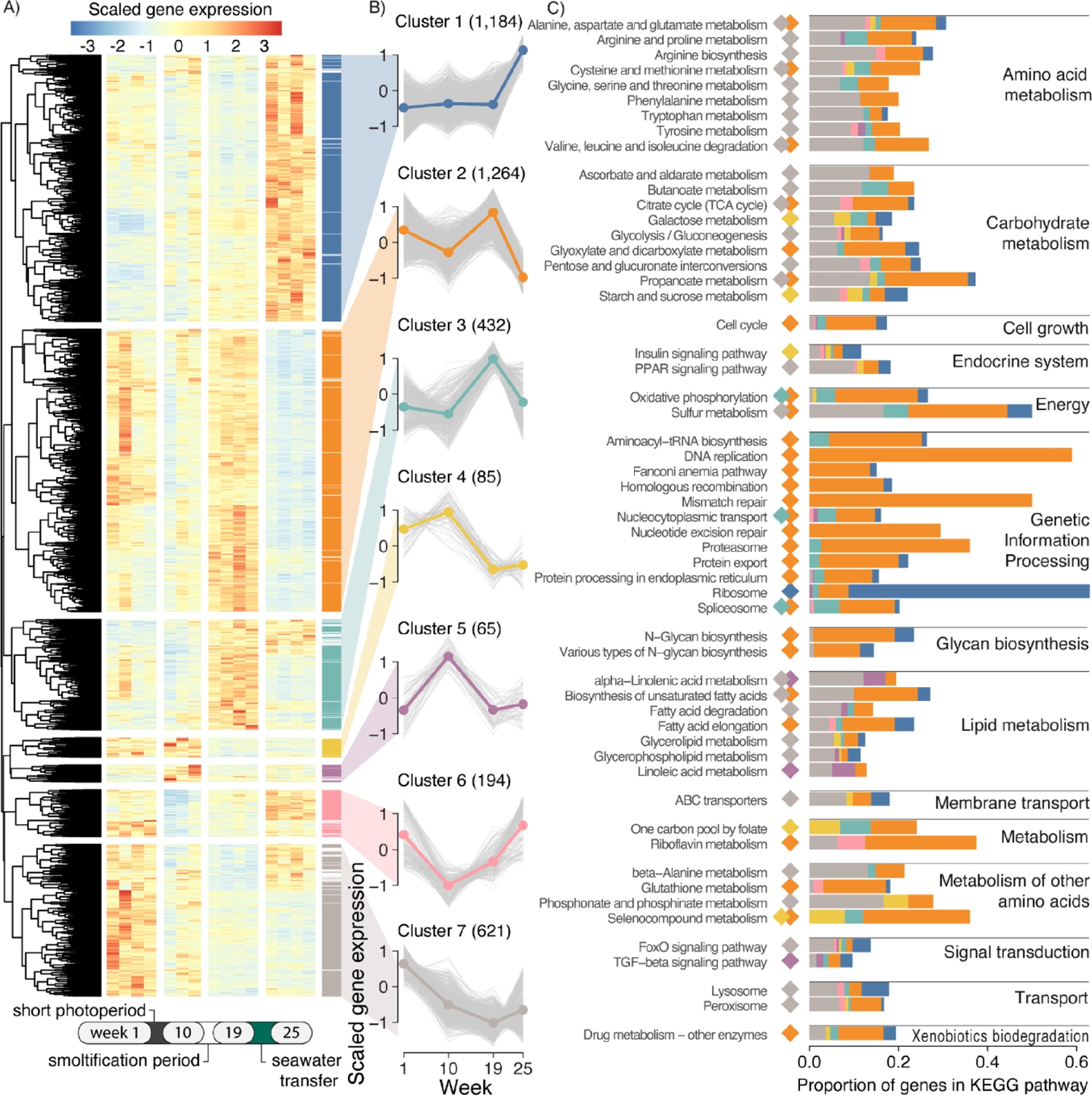
Global gene expression changes across life-stage. A) Relative liver expression of genes differentially expressed between any time point in the experimental fish cohort (FDR <0.05). Scaled expression is denoted as gene-scaled transcripts per million. Genes were partitioned into six co-expression clusters by hierarchical clustering. Colored bars indicate cluster membership when correlation to the mean cluster pattern was >0.5. Genes with correlation =<0.5 were excluded. B) Gene expression trends over time by cluster. Colored line indicates mean relative expression while grey lines are individual genes within the cluster. C) KEGG pathway enrichment by cluster. Colored diamonds indicate for pathways which clusters they are significantly enriched in (adjusted p <0.05). Colored bars indicate the proportion of genes within the pathways that are in clusters.

To associate well defined metabolic or signaling processes to the different gene expression trends, we performed KEGG enrichment analysis on each co-expression cluster, yielding 56 unique significantly enriched (adjusted p <0.05) pathways (Fig 2C). Genes in clusters 2 and 3 that increased during smoltification and sharply decreased after seawater transfer were enriched in pathways related to genetic information processing, cell growth, protein metabolism, and oxidative phosphorylation. Genes in clusters 4 and 5 which had peak expression after a short photoperiod and decreased during smoltification and seawater transfer were similarly enriched in genetic information processing pathways and energy metabolism, however they also contained several pathways related to amino acid metabolism including cysteine and methionine metabolism, glutathione metabolism, and selenocompound metabolism. Cluster 1 genes strongly increased in relative expression after seawater transfer and was exclusively enriched in the ribosome pathway. Genes in cluster 7 which decreased in relative expression during smoltification and remained low during seawater transfer were enriched mainly in lipid, amino acid, and carbohydrate metabolic pathways, ABC transporters, and signaling pathways including FoxO signaling and PPAR signaling.

Since many KEGG pathways contain enzymes with reciprocal activities, we manually examined genes within select enriched KEGG pathways to determine what was driving enrichment trends. In lipid metabolic pathways we observed a distinct bias in genes relating to long-chain fatty acids towards downregulation in freshwater smolts. Seven long-chain-fatty-acyl-CoA ligase (*acsl*) genes (*acsl1*, three *acsl3* and three *acsl4*), acetyl-CoA carboxylase (*acc1*), three acetyl-CoA synthetase genes (*acs2l-1*, *acs2l-1*, and *acs2l-1*), several key genes related to polyunsaturated fatty acid (PUFA) biosynthesis (*5fad*, *6fada*, *6fadb*, *elovl5b*, and *elovl2*), both copies of fatty acid synthase (*fas1* and *fas2*), and three copies of the key gene diacylglycerol acetyltransferase (two *dgat1* and one *dgat2*) all significantly decreased during smoltification and remain lowly expressed through seawater transfer (Fig 3B).

**Fig 3:**
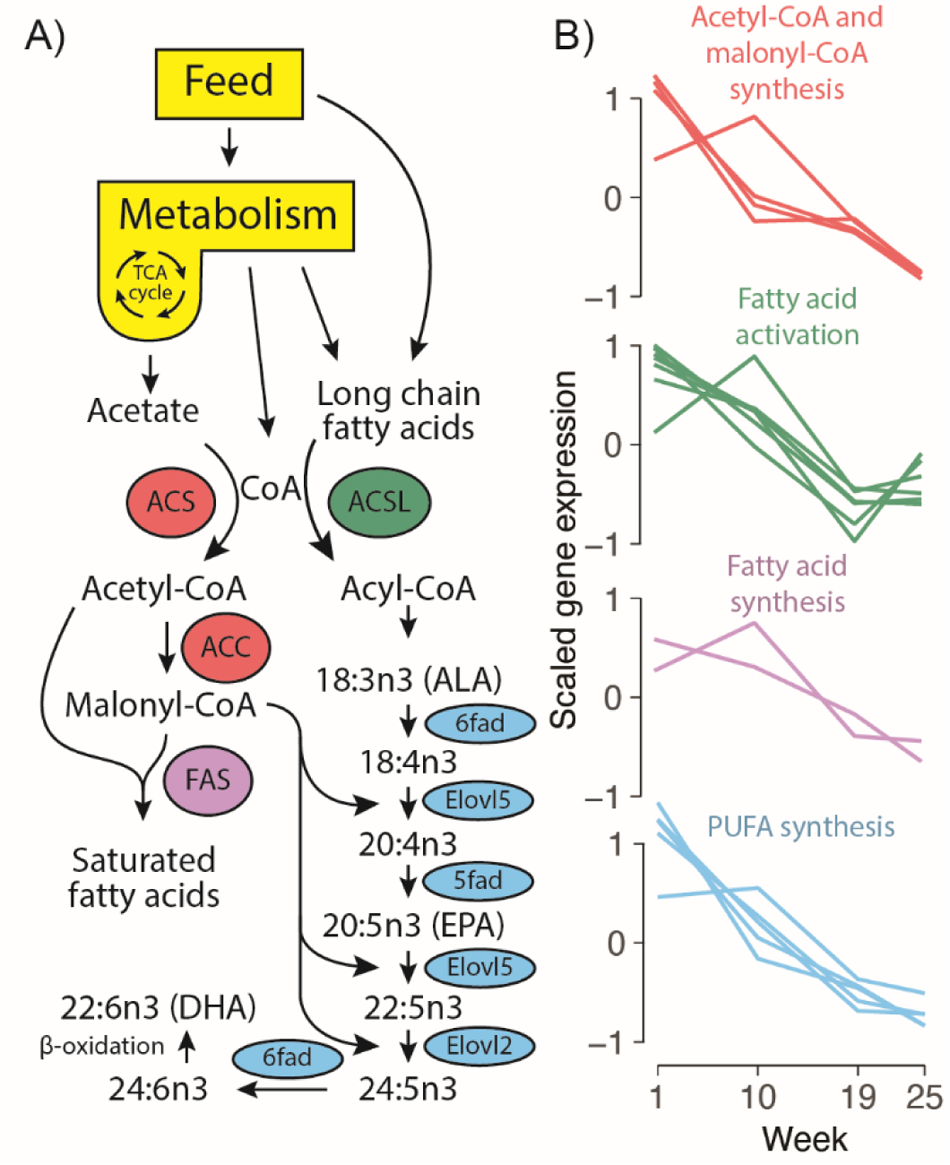
Expression of key lipid metabolism genes across the parr-smolt transition. A) Schematic of the lipid biosynthesis pathway in Atlantic salmon. B) Relative expression of genes in the pathway over time. Acetyl-CoA and malonyl-CoA synthesis (red) displays genes *acc1*, *acs2l-1*, *acs2l-2*, and *acs2l-3*. Fatty acid activation (green) displays genes *acsl1*, *acsl3l-1*, *acsl3l-2*, *acsl3l-3*, *acsl4*, *acsl4l-1*, *acsl4l-2*. Fatty acid synthesis (purple) displays gene *fas1* and *fas2*. Poly unsaturated fatty acid (PUFA) synthesis (blue) displays genes *5fad*, *6fada*, *6fadb*, *elovl2*, and *elovl5b*.

Synthesis of acetyl-CoA by *acs* and activation of long-chain fatty acids by *acsl* is the first obligatory step for entry into beta-oxidation or biosynthesis pathways (Fig 3A), so a decrease in these gene products likely means that metabolism of their substrates (acetate and C12 to C20 fatty acids) also decreases [15]. Finally, three copies of the key gene diacylglycerol acetyltransferase (two *dgat1* and one *dgat2*) which catalyzes the last committed step in triacylglycerol biosynthesis [16] decreased in freshwater smolts. Collectively, co-downregulation of these important lipid associated genes is a strong indicator of decreased utilization and processing of fatty acids, especially long chain poly unsaturated fatty acids (LC-PUFA), in smolts preparing to enter a seawater environment.

We identified genes directly influenced by seawater transfer by performing a pair-wise test for gene expression changes between the experimental group and freshwater control group at week 25. This resulted in 2,121 DEGs (FDR <0.05, Table S5), most (1227) being downregulated in seawater compared to freshwater (Fig S1) and overlapped with many genes belonging to co-expression clusters 1 (281) and cluster 2 (529) in Fig 2. Regarding lipid metabolism, processes related to de novo fatty acid synthesis decrease in seawater relative to control. Both copies of fatty acid synthase (*fas1* and *fas2*) and one acetyl-CoA carboxylase (*acc1*) decreased expression in seawater, all of which catalyze key steps in *de novo* fatty acid synthesis [17]. Additionally, two other *acsl* genes (*acsbg2* and *acsl1*) known to be involved in saturated and monounsaturated fatty acid activation were downregulated in seawater [18]. This coincided with an increase in several thioesterase genes, including *acot1l* and *acot5l*, responsible for de-activation of fatty acids through the hydrolysis of acyl-CoAs [19]. It is unlikely that de-smoltification occurred in the freshwater control smolts because expression of these genes remains stable between weeks 19 and 25 in the control fish. This combination of decreased expression of key *de novo* biosynthesis genes and increased fatty acid de-activation through greatly increased thioesterase expression suggests a reduced capacity to synthesize fatty acids in liver of fish after transition to sea, in line with previous findings [14].

### No long-term effect of short photoperiod exposure in liver

To evaluate the role of photoperiodic history on the development of liver function during smoltification, we performed pair-wise tests for gene expression changes between experimental fish exposed to a short photoperiod and control fish on constant light regime at week 10 (at the end of the short photoperiod exposure) and at week 19 (just prior to seawater transfer). We identified a relatively shorter list of DEGs (532, FDR <0.05, Table S3) associated with photoperiodic history differences at week 10, but only a few DEGs at week 19 (15, Fig 4, Table S4). At week 10 we found a vitamin D 25-hydroxylase gene, the first step in the formation of biologically active vitamin D [20], was strongly downregulated after a short photoperiod. This is likely due to decreased UV mediated vitamin D synthesis in the skin from less exposure to light [21]. The low number of DEGs at week 19, and low overlap between week 10 and 19 expression changes, showed that different photoperiodic histories did not impact longer term gene function in the liver following smoltification.

**Fig 4:**
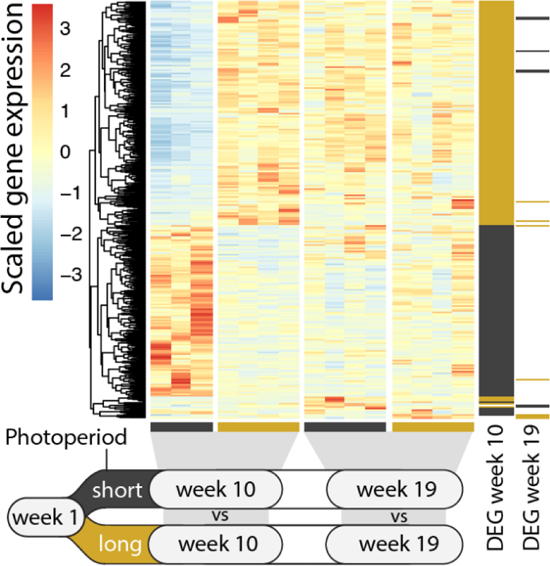
Gene expression changes in response to photoperiod history. Relative liver expression of differentially expressed genes (DEGs, FDR <0.05) between the short photoperiod exposed experimental group and long photoperiod control group at weeks 10 and 19. Genes are marked to the right if differentially expressed higher (black) or lower (yellow) in response to a short photoperiod at weeks 10 and/or 19.

### Remodeling of liver transcription factor binding across smoltification

To better understand the mechanistic drivers shaping changes in liver gene expression through salmon life stages, we generated ATAC-seq data to measure accessibility of chromatin and used this to indirectly quantify transcription factor (TF) occupancy at predicted TF binding sites (TFBS) by assessing local drops in chromatin accessibility (aka footprints). For each time point across the 25 week experiment group we generated two replicate samples of ATAC-seq data from the same livers sampled for RNA-seq, at a depth of 55-72M reads. Reads were aligned to the genome (41-63M) and peaks where reads were concentrated were called to represent regions of accessible chromatin. A unified set of the ATAC peaks was made by merging peaks across the different weeks (File S1). A principal component analysis (PCA) on the sample’s read counts over the unified peaks showed pairing of replicates and separation between the weeks (Fig 5A). PC1 separated weeks 1 and 19 to 10 and 25, while PC2 separated the pre-smolt weeks (1 and 10) to the post-smolt (19 and 25). The unified set of 201k peaks was composed of peak sets from each week, with week 19 having the highest number of peaks (181k, Fig 5B). Most of the peaks at each week were shared across sets, with week 19 standing out as having the greatest number of unique peaks (Fig 5C). Peaks were highly concentrated around the TSS of genes as expected, associating with gene regulation (Fig 5D). Peaks were mostly found in introns or intergenic regions, suggesting a higher proportion of peaks at enhancer than promoter elements, not unexpected (Fig 5E).

**Fig 5:**
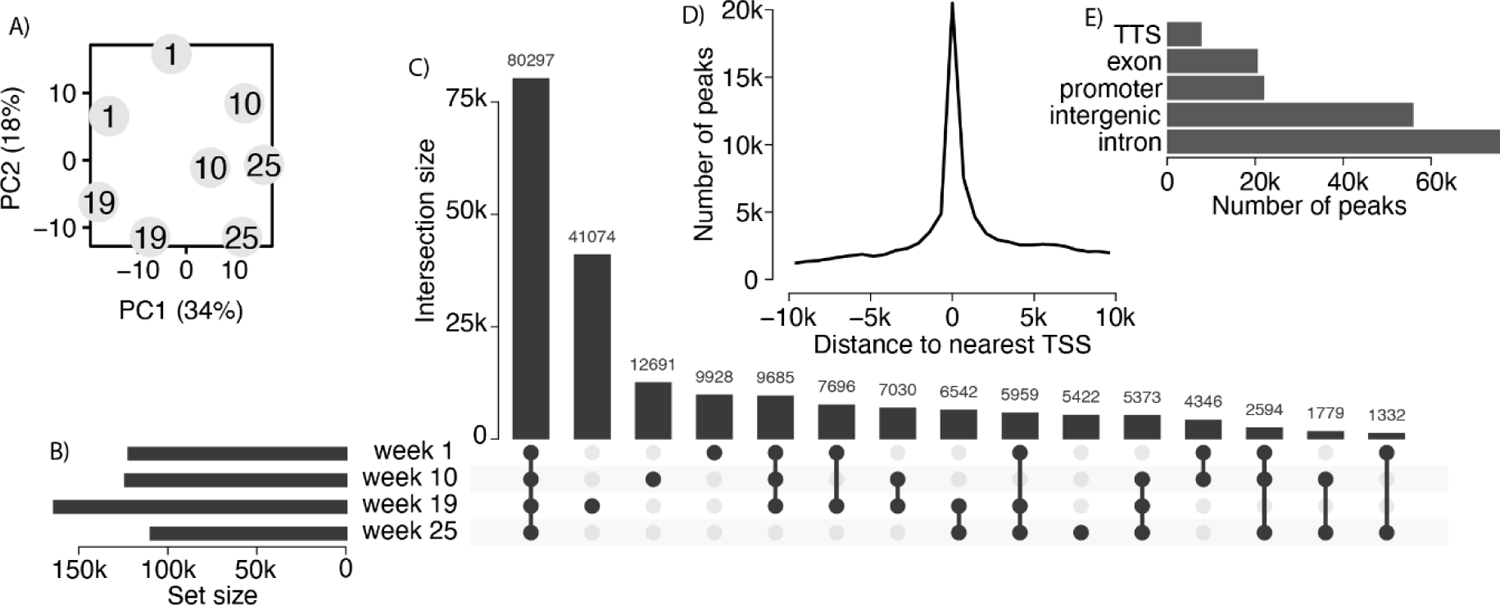
Comparison of chromatin accessibility across life-stage. A) Similarity of ATAC-seq samples at different weeks by principal component (PC) analysis of ATAC-seq read counts within a unified set of ATAC peaks (shared and unique between weeks). B) Number of ATAC peaks called at each week. C) Number of peaks intersecting between sets or unique to each week. D) Distribution of distances (in base pairs) of unified peaks to the nearest gene transcription start site (TSS). E) Genomic locations of unified peaks.

Little is known about the environmentally driven changes in gene regulatory pathways in salmon. We therefore used a TF footprinting analysis to identify within peaks drops in reads at TFBS, indicating a bound TF (i.e. occupancy) at that site in the given sample. We first describe the TFs showing genome-wide changes in TFBS binding between livers sampled in different photoperiods and water salinities (Table S6). Since developmental stage (age) can also impact TF binding patterns, we focused on TFs with differences in occupancy that persists across environmental contrasts with fish from different developmental stages (Fig 6A and B). Using a cutoff for differential TF occupancy (log_2_ fold change in genome wide TF-motif occupancy >0.1) we identified 33 and 35 TF binding motifs that were associated with photoperiod and salinity, respectively (Fig 6C, Table S7).

**Fig 6:**
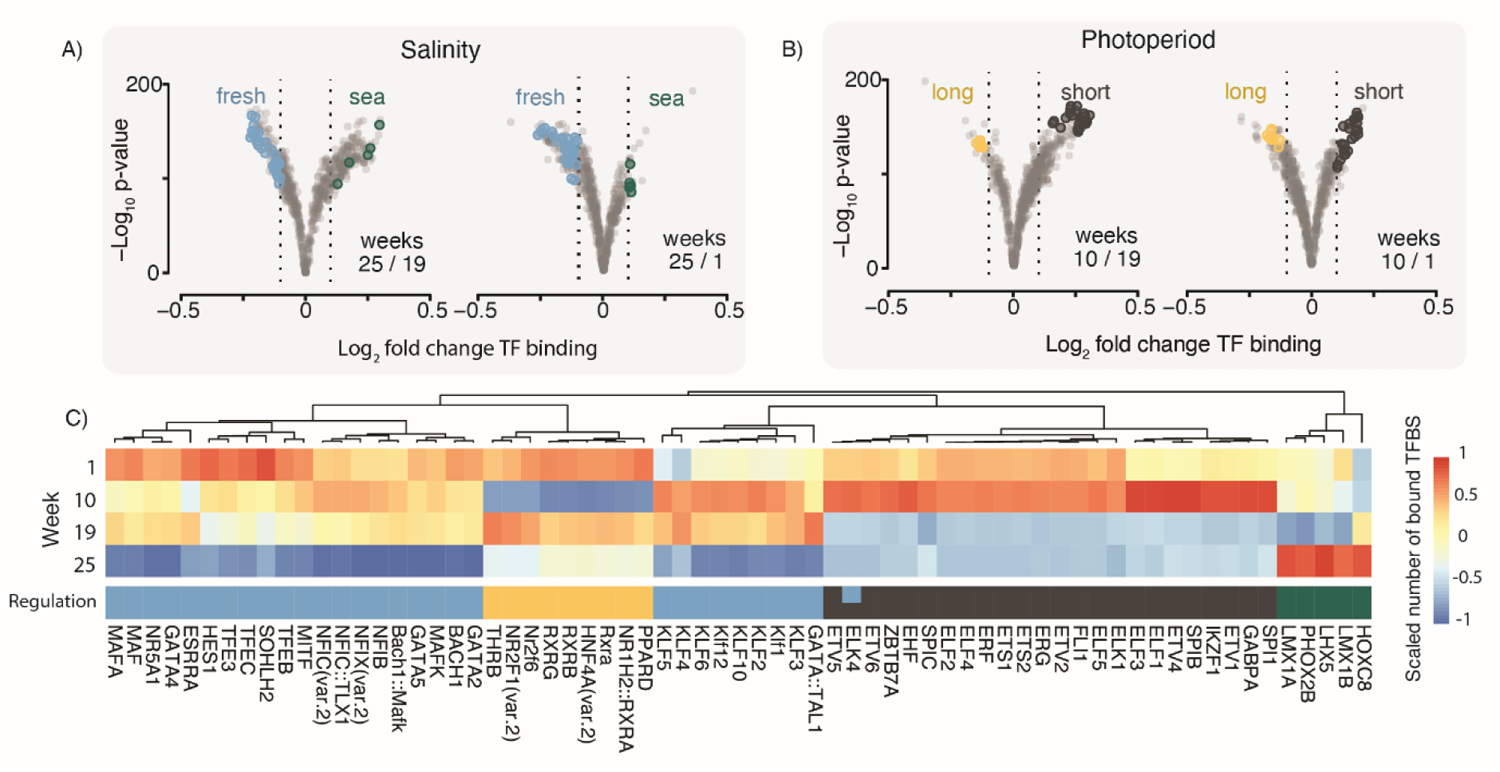
Genome wide changes in transcription factor activities across key life stages. Volcano plots show genome wide fold changes and significance for transcription factor binding site (TFBS) binding scores between A) fresh- to saltwater weeks and B) short to long photoperiod weeks. Transcription factor (TF) motifs with significant changes in global binding scores (absolute log_2_ fold change >0.1) across each contrast are colored. C) Heatmap shows the scaled number of total bound TFBS across the weeks for the TF motifs that are significant in A) and B). The ‘regulation’ color indicates in which environment the TF motif had the greater TFBS binding score.

Most (30/35) TF motifs associated with water salinity differences were found to have a marked drop in genomic occupancy after transition to saltwater (Fig 6C). These include several motifs known to bind TFs associated in energy homeostasis related processes, such as TEF’s, GATA4, NR5A1 MAF, KLFs [22–24]. Only five TF motifs had a significantly higher occupancy in saltwater, including LIM’s and two homeobox binding motifs (PHOX2B and HOXC8). The photoperiod contrast (Fig 6B and 6C) revealed that most TFBS’s with induced occupancy after a short day period were binding sites for E-26 family transcription factors (ETS, ERG, ETV, SPIC, ELK, SPI1), which have been associated with regulation of circadian genes in other species [25, 26]. It is interesting to note that these TF binding sites with a spike in occupancy after a short photoperiod drops dramatically towards the end of smoltification (week 19) and stays low after seawater transfer (week 25). TF binding sites with reduced occupancy following a period of shorter days were mostly related to homeostasis of cellular metabolism, including key liver glucose and lipid metabolism regulators PPARD [27], RXRs [28] and HNF4A [29], as well as occupancy of thyroid hormone receptor beta (THRB) [30].

Next, we wanted to link TF binding patterns to the specific gene regulatory dynamics. We assigned TF motifs to genes (i.e. motif-gene pairs) by closest proximity and asked if genes with a particular expression pattern (Fig 7A) were enriched for TF motifs in proximity with a corresponding pattern of TF occupancy (Fig 7B). For example, for genes in expression cluster 1 with highest expression at week 25, we expect nearby binding sites to be enriched in TF motifs that are occupied by TFs in week 25, but not the other time points. Indeed, genes in most expression clusters displayed significant enrichments of TF motifs with expected binding patterns (colored red or orange), and these signatures were quite distinct for each gene expression cluster (Fig 7C, Table S8). A diagram shows the comparison for the fisher’s exact tests between gene expression cluster and TF binding times to find TF motif significance (Fig 7D).

**Fig 7.**
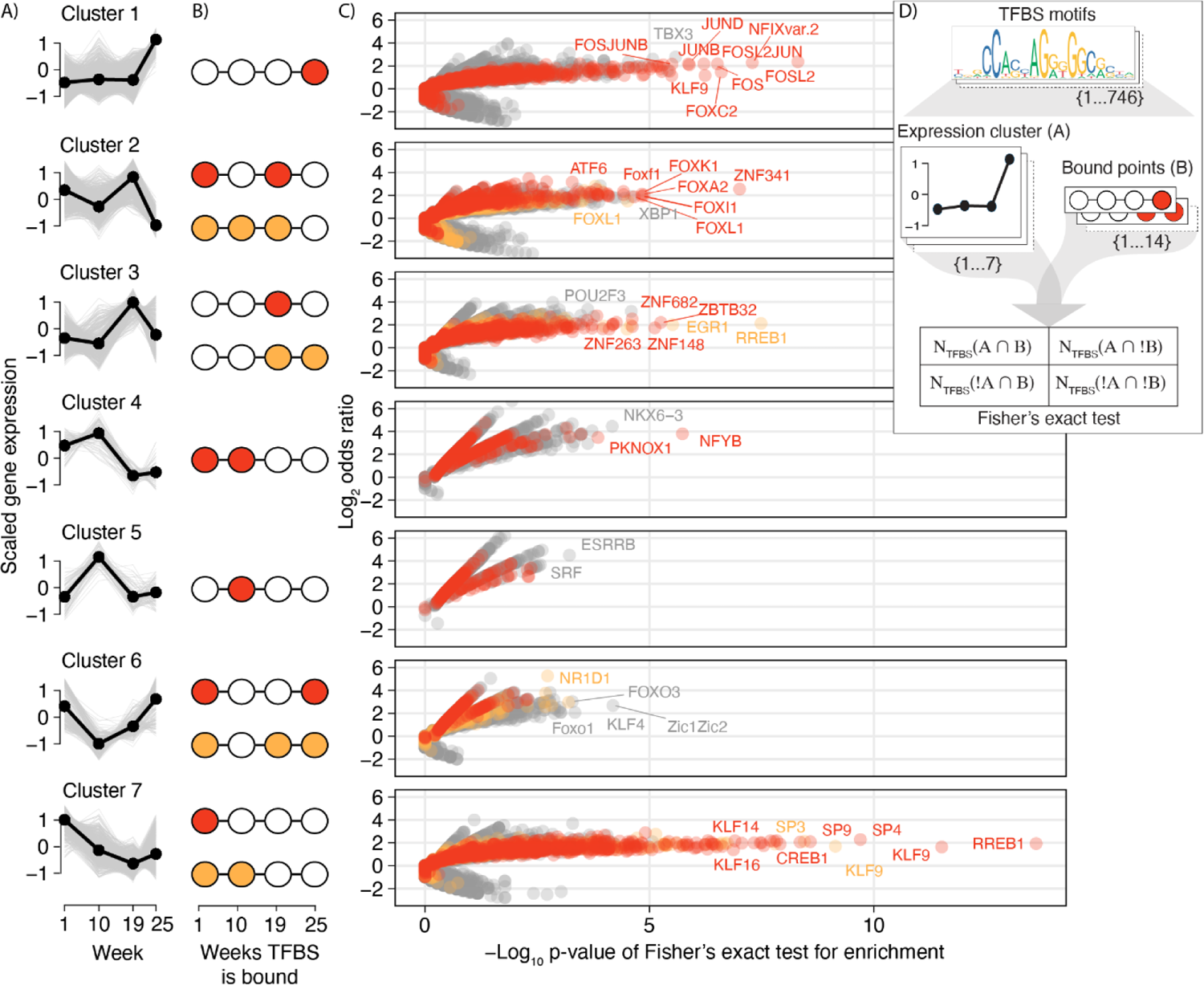
Enrichment of transcription factor binding patterns within gene expression clusters. A) Gene expression trends of clusters from Fig 1. B) Assumed weeks the transcription factor binding sites (TFBS) near genes in the cluster would be bound by a transcription factor (TF) to regulate transcription at the weeks of highest expression. A primary and secondary assumed binding pattern is colored red and orange respectively. C) For each set of genes in a cluster, each TF was tested if the TFBS near to those genes were enriched for any combination of binding pattern. Fisher’s exact test results for all TF-binding pattern combinations are plotted per cluster as odds ratios against the significance. The test results for the assumed primary and secondary binding patterns in B) are colored red and orange, respectively. A top proportion of the most significant TFs (in a top quantile per test) are labeled. D) Diagram showing how each combination of TFBS motif, expression cluster, and binding pattern was tested for enrichment with a Fisher’s exact test.

From the enrichment results (Fig 7C) we see genes in cluster 1, enriched for ribosome related functions, had nearby TFBS with NFIX binding following transition to seawater. This is a transcriptional regulator known to be involved in ribosome biogenesis [31]. In addition, cluster 1 TFBS were associated with binding of FOS and JUN after seawater entry. These TFs are major components of the Activator Protein 1 (AP-1) transcription factor complex which is responsive to growth factors and drive cell proliferation and differentiation [32–34]. The top associated TFs for cluster 2 genes were ZNF341 known to be involved in immune homeostasis [35] and several Fox TFs (A, F, L, I, K) linked to various cell physiological processes [36, 37]. Cluster 3 genes were associated with several unnamed zinc finger transcription factors (ZNFs and ZBTBs) binding at week 19 prior to seawater transfer, as well as binding of RREB1 and EGR1 in week 19 and week 25. Among TFs with binding in week 19 only, we find genes linked to regulation of immune cell function (ZBTB32 and ZNF263) [38, 39] as well as oxidative phosphorylation [40]. The RREB1 and EGR1 are well described players in RAS signaling pathways [41–43] involved in cell growth and proliferation. Among cluster 4 genes we find enrichment of insulin and sugar metabolism functions, with PKNOX1 and NFYB TF binding significantly associated with their expression patterns. Both these TFs have been shown to function in lipid metabolism and be linked to insulin signaling [44–46]. The top TFs associated with cluster 6 gene expression is NR1D1 (also called Reverbα), a core component of the circadian clock and regulator of lipid metabolism [47]. Cluster 6 is not enriched for any KEGG pathways but has a marked drop in expression after the short photoperiod exposure. In the final cluster 7, enriched for genes playing roles in amino acid, glucose, and lipid metabolism, we find very strong associations with binding of several TFs, including KLF and SP family members. Indeed, these TFs are known to play important roles in regulating gluconeogenesis and lipid and amino acid metabolism in mammalian livers [24, 48].

### DNA methylation not linked with gene expression

To investigate the role DNA methylation had on the gene expression changes throughout salmon life-stages, we produced a RRBS dataset from the same liver samples at a depth of 26-40 M reads. A consensus set of 1.2M CpG positions was used for differential methylation analysis (Table S9). To assess genome wide differences in the regulation of CpG methylation, a principal component analysis (PCA) separated samples based on the methylation levels across the CpG consensus set. There was no clear separation of samples by timepoint or experiential condition, with PC1 and PC2 each explaining less than 5% of the variance (Fig S3A). To find specific sites of differential methylation, we tested all CpGs for differences in methylation score across any timepoints of the experimental group with an ANOVA-like approach (Table S10). Out of the consensus set of CpGs, 2535 (0.2%) were differentially methylated cytosines (DMC) across life-stages (FDR <0.05, fold change >25%, Fig S3D-F). 209 of these were present in promoter regions, 103 in exons, 664 in introns, and 782 in intergenic regions (Fig S3E). Most genomic regions with a DMC had one differentially methylated site and only a few regions contained longer stretches (up to 26 bp in an intron) of differentially methylated CpGs (Fig 3D). We assessed if these CpGs were associated to genes with a specific function, but enrichment tests for GO or KEGG terms gave no significant results. We looked at next if the methylation percentages of DMCs correlated to changes in their corresponding gene’s expression across timepoints. 157 DMC-gene pairs were significantly correlated (p <0.05, Pearson correlation coefficient >0.95), however most of these genes were relatively lowly expressed. Simulating random DMC-gene pair correlation values found that the distribution of values from our real data was not significantly different from that of simulated pairs (p-value <0.62, Fig S3G), refuting strong links between regulation of CpG methylation and gene expression in our experiment.

## Discussion

### Lipid metabolism remodeling as a pre-adaptation to life at sea

An important feature of smoltification is how the process prepares the juvenile fish for a life in sea, i.e. physiological necessities for survival are already present while the fish is still in freshwater. This is well documented for the osmoregulatory machinery in the gills [49–54]. For example, the likely causal agent for saltwater tolerance, NKA a1b, increases in abundance in freshwater gill tissues [8]. While lipid metabolism related gene expression is known to decrease in liver of seawater stage Atlantic salmon [14], this is the first report that systemic downregulation of lipid metabolism gene expression actually occurs before transition to sea (Figs 2 and 3). Given the availability of polyunsaturated fatty acids in seawater environments is higher than freshwater [55], and that the body lipid composition changes to match this in freshwater smolts [56], it is likely that the observed decrease in lipid metabolism is a genetically programmed preadaptation to life at sea. This study therefore adds another feature to the list of pre-adaptations in freshwater smolts.

### The effect of photoperiod history on genome regulation during smolt liver development

Decades of research have revealed that many features of smolt physiology development can be affected by photoperiodic history [3, 57–64], including gene expression patterns. For example, a recent study of gill transcriptomes identified a subset of 96 genes with significantly increased gene expression levels in smolt exposed to a short photoperiod (8:16) during development compared to smolts kept on constant light (24:0) [3]. In this study, however, no long-term effects of short photoperiod exposure were found in liver transcriptomes (Fig 4). We do, however, show that salmon liver gene regulation is responsive to variation in photoperiods. Transcriptome profiles through smolt development show distinct gene sets with increased and decreased expression after reduction in photoperiod (Fig 2A clusters 5 and 6). Furthermore, analyses of TF binding dynamics (Fig 6) identify photoperiod sensitive TFs encoded by genes known to have repressed expression under short photoperiod in mammals, such as retinoic acid related TFs (RXRs) and thyroid hormone receptors (THRB) [65, 66]. In addition, our integrated analyses of gene expression profiles and TF binding (Fig 7) associated NR1D1, a core component of the mammalian circadian clock [47], with genes having lower expression after exposure to short photoperiod. Taken together, we conclude that smolt liver development does not seem to rely on having experienced a winter-like photoperiod. Yet since acute effects of reduced photoperiods had a large impact on gene regulatory networks related to metabolism (Figs 6 and 7), it is likely that highly divergent photoperiodic histories can lead to delayed spillover effects and result in differences in metabolic states.

### Linking genome regulatory layers to understand the developing smolt liver

Salmon experience gene regulatory changes [13, 14] and function [6, 7] in liver during smolt development in freshwater, and following seawater entry. Yet, no genome wide studies of DNA methylation and TFs involved in driving these transcriptional and physiological changes have been conducted. Here, we generated an RRBS dataset as well as an ATAC-seq dataset across smolt development in liver and used the latter to predict TF occupancy dynamics and map out putative major regulators of key developmental processes during smoltification (Figs 6 and 7).

Firstly, we showed that dynamic DNA methylation has a limited role in gene regulation in the liver during smoltification. Of the 1.17 M CpGs in our dataset, only about 2500 of these showed dynamic methylation during smoltification, and few of these were in the vicinity of differentially expressed genes. This echoes an earlier study on methylation changes associated with early maturation in Atlantic salmon in which the liver exhibits less dynamic methylation overall than brain and gonads [67]. Also in other organs of Atlantic salmon, gene expression changes seem to be controlled by other gene regulatory features than DNA methylation [68]. Despite the intriguing hypothesis of DNA methylation being an important gene expression regulator, and an epigenetic one at that, our study questions this role in the context of post-embryonic development. Indeed, many studies describe methylome changes during metamorphosis or other post-embryonic transitions but does not provide strong evidence for a causative connection between changes in gene expression and changes in DNA methylation [69–71].

Previous studies of liver physiology during parr-smolt transformation highlights decreased lipid and glycogen biosynthesis and increased levels of glyco- and lipolysis [7]. In line with this, about 600 genes enriched for lipid, carbohydrate, and amino acid metabolism related functions displayed a clear decreasing trend in gene expression from parr (week 1) to smolt (week 19) (Fig 2, cluster 7). Several TFs showed highly significant TFBS binding associations with the cluster 7 gene expression profile (Fig 7C) including KLF/SP gene family members known to play important roles in regulating gluconeogenesis and lipid and amino acid metabolism in mammalian livers [24, 48]. Finally, genes in expression cluster 4, also showing a marked decrease in expression in smolts (week 19), were enriched for TFBS that had a significant drop in NFY binding from week 19. This TF is known to be a major regulator of lipid metabolism, including biosynthesis of fatty acids [44]. Concurrently, but with opposite expression trends, genes involved in oxidative phosphorylation related functions (the last step in breaking down amino acids, lipids, and carbohydrates to energy) increased in expression from parr to smolts (Fig 2, cluster 3). The TFBS of these same genes were enriched for binding of the TF ZNF682 in smolts in week 19 (Fig 7C), a nuclear encoded TF gene that regulates oxidative phosphorylation in human cells [40]. Together, these results suggest that increased ZNF682 occupancy in combination with reduced KLF and NFY promoter binding has an associated link to the liver metabolic shift from synthesis to break down of organic compounds as fish undergo parr-smolt transformation.

Following the parr-smolt transformation, the transition to a life in seawater is also known to be associated with additional changes in physiology in Atlantic salmon. Genes increasing in expression in seawater (Fig 2, cluster 1) were involved in ribosome biogenesis and their TFBS were associated with increased NFIX occupancy in seawater (Fig 7C), reported to impact ribosome biogenesis in mammals [31]. Another well known route to increased ribosome gene expression and protein synthesis is the induction of the mTOR pathway [72, 73]. Interestingly, seawater entry is known to trigger increased growth hormone levels in salmon [74, 75] and this hormone acts as a rapid activator of protein synthesis through the mTOR pathway [72]. Furthermore, seawater growth hormone increase can also be linked to the second group of TFs putatively involved in gene expression induction after seawater transition (Fig 2), namely JUN and FOS (Fig 7C). These genes, originally known as onco-genes, are also responsive to growth hormones, and regulate cell proliferation and differentiation [32–34]. These TFs provide the molecular basis for linking growth hormones to increased growth capacity of smolt in seawater [4]. In our experiment freshwater control fish were larger than fish transferred to sea, but this can be explained by a known initial suppression in growth and feeding followed by increased growth rates [76]. Finally, genome-wide footprint signals (Fig 6) also pointed to large changes in the binding of TFs involved in energy homeostasis after seawater entry [22–24], further underpinning the metabolic gear shift. In conclusion, our data suggested that the genome regulatory dynamics in smolt livers across the fresh- to seawater transition is likely driven to a large extent by an increase in circulating growth hormone, resulting in activation of major regulatory pathways (e.g. JUN/FOS) for cell growth and differentiation.

## Materials and methods

### Smoltification trial

Atlantic salmon eggs, provided by AquaGen Breeding Centre Kyrksæterøra, Norway, were sterilized at the Norwegian University of Life Sciences (NMBU) fish lab and incubated at 350 to 372 day-degrees until hatching. First feeding of fry was five weeks after hatching when the egg sac had been depleted. Fry were reared in two replicate tanks and on a standard commercial diet high in EPA and DHA fats for the duration of the trial. Fish occasionally needed to be euthanized as they grew to maintain adequate dissolved oxygen levels in the tanks. Sampling began 21 weeks after first feeding as week 1, and again at weeks 10, 19, and 25. Sampled fish were euthanized by a blow to the head and samples of liver tissue were cut into ∼5 mm cubes, placed in RNAlater, and incubated for at least 30 minutes at room temperature before long-term storage at −20°C. One week after the first sampling, some fish from each tank were transferred to replicate photoperiod control tanks where the day length remained unchanged. At the same time, the experimental tanks’ photoperiod was switched to “winter-like” lighting conditions with 8 hours of light per day for 8 weeks to trigger smoltification before returning to “spring-like” conditions with 24 hours of light per day. Immediately after the week 19 sampling, some fish from each experimental tank were transferred to seawater conditions at the Norwegian Institute for Water Research (NIVA), Solbergstranda, Norway. UV-sterilized seawater used in this life-stage had a salinity of 3%-3.5% and was obtained from the Oslofjord. Fish were sedated before transport and allowed to acclimatize for several hours before being slowly introduced to the new water conditions. The fish that were not transferred to seawater were sampled as freshwater controls at the same time as the experimental fish. All animals used in this study were handled in accordance with the Norwegian Animal Welfare Act of 19th June 2009.

### RNA sequencing

For RNA sequencing we extracted total RNA of liver samples from experimental and control groups taken on weeks 1, 10, 19, and 25 in replicates of four with the RNeasy Plus Universal Kit (QIAGEN). Concentration was determined with a nanodrop 8000 spectrophotometer (Thermo Scientific) and quality was assessed by running on a 2100 bioanalyzer using the RNA 6000 Nano Kit (Agilent). Extracted RNA with an RNA integrity number (RIN) of at least eight was used to make RNA-seq libraries using the TruSeq Stranded mRNA HT Sample Prep Kit (Illumina). Mean length and library concentration was determined by running libraries on a 2100 bioanalyzer using a DNA 1000 Kit (Agilent). RNA-seq libraries were sequenced by the Norwegian Sequencing Center (Oslo, Norway) on an Illumina HiSeq 4000 using 100 bp single end reads and at a depth of 25-43M reads per sample.

### Gene expression quantification

Gene expression was quantified from RNA-seq fastq files through the nf-core rnaseq pipeline (v3.9), which involves quality control, read trimming and filtering, read alignments to the Atlantic salmon genome and gene annotations (NCBI refseq 100: GCF_000233375.1_ICSASG_v2) with STAR aligner, and read quantification from alignment with the salmon program. See pipeline documentation for further details on all steps: nf-co.re/rnaseq. Gene level counts, length scaled, were used for differential expression testing, and gene transcript per million (TPM) values used for visualizations including expression heatmaps and line plots. A PCA of gene TPMs showed one sample at week 10 (week_10_2_3, Biosample: SAMEA14383461) as an outlier (Fig S2) so we removed this sample from the analysis.

### Differential expression analysis

Differentially expressed genes (DEGs) were tested for first differences across all experimental group time points using an ANOVA-like test with edgeR R package (v3.36) [77], using the generalized linear model and quasi-likelihood F-test function (glmQLFTest), testing for differences between weeks 1, 10, 19, and 25. DEGs were chosen from the results using an FDR cutoff of <0.05. Euclidean distances of DEG were calculated based on TPM values over the samples, and the DEGs separated into 7 clusters. We chose 7 clusters based on observing patterns in the heatmap of expression and comparing the sum of squares within and between different numbers of clusters. DEGs had to have correlation >0.5 to the mean expression values of their assigned cluster, otherwise they were excluded from the cluster. DEGs were also tested between experimental and control groups (short and long photoperiod history, respectively) at week 10 and 19 separately, in pair-wise exact tests with edgeR (v3.36), choosing DEGs with an FDR cutoff of <0.05. Enrichment of KEGG pathways in sets of DEGs was performed with the clusterProfiler R package (v4.2.2), using the pathway data for Atlantic salmon genes within the KEGG database.

### ATAC sequencing

The protocol for the ATAC assay was based on that in Buenrostro et al. 2013 [78]. Two replicate liver tissue samples were used from the previous sampling of the experimental group on weeks 1, 10, 19 and 25. The liver tissues were washed and perfused with cold PBS to remove blood before being dissociated and strained through a cell strainer. The nuclei were isolated from cell homogenate by centrifugation and counted on an automated cell counter (TC20 BioRad, range 4-6 um). Transposition of 100k (weeks 1, 19 and 25) and 75k (week 10) nuclei was performed by Tn5 transposase from Nextera DNA Library Preparation kit. The resulting DNA fragments were purified and stored at −20°C. PCR Amplification with addition of sequencing indexes (Nextera DNA CD) were done according to Buenrostro et al. 2015 [79], with a test PCR performed to determine the correct number of amplification cycles. The ATAC libraries were cleaned by Ampure XP beads and assessed by BioAnalyser (Agilent) using High sensitivity chips. Quantity of libraries were determined by using Qubit Fluorometer (Thermo). Mean insert size for the libraries was 190 bp. Sequencing was done on a HiSeqX lane using 150 bp paired-end reads and at a depth of 55-72M reads per sample.

### ATAC peak calling

Calling of ATAC-seq peaks was done through the nf-core atacseq pipeline (v1.2.1), which involves quality control, read trimming and filtering, read alignments to the Atlantic salmon genome (BWA aligner), and calling of narrow peaks per sample as well as a unified narrow peak set across all samples (MACS2 peak-caller). Data for QC results including PCA of samples, intersection of peaks sets, peak distances to TSS, and peak genomic locations, were also computed through the pipeline. See pipeline documentation for further details on all steps: nf-co.re/atacseq.

### TF footprinting

TF footing in the unified ATAC peak set previously generated was done using the TOBIAS program (v7.14.0) [80]. With it, we identified within peaks using the ATAC-seq read alignment BAM files for each week (reads from replicates combined) ‘footprints’ (i.e. dips in read depth within peaks), indicative of TF proteins binding to the DNA and locally blocking transposase activity during the ATAC protocol. We used a set of TFBS motifs from the JASPER database (2020 CORE vertebrates non-redundant) to associate these footprints with specific TFs. Peaks were associated to a gene by closest proximity to the TSS during the atacseq pipeline. A blacklist file was used to mask simple repetitive regions in the genome from analysis (generated in-house). Data for the genome-wide changes in TF binding was taken from the ‘bindetect_results.txt’ file produced by TOBIAS, plotting the change in TF binding scores between weeks against their p-values. The number of bound sites for each TF was scaled across weeks and used for the heatmap visualization of differences. We identified TFs changing in response to salinity or photoperiod by intersecting TF binding results between the different weeks. With a log_2_ fold change cutoff of >0.1, TFs that had an increase in binding in week 25 compared to both weeks 1 and 19 were assigned as ‘sea’, and inversely those with a decrease assigned as ‘fresh’. Similarly for photoperiod those increasing in week 10 compared to 1 and 19 were assigned ‘short’ and those decreasing were assigned ‘long’. We tested for the enrichment of certain TF binding patterns (which weeks TFBS were bound) within the gene expression clusters previously identified. For this test we used the peak-gene annotation data mentioned previously to assign the TFBS to genes. Then for each combination of TFBS motif type, expression cluster, and TF binding pattern, we used a Fisher’s exact test to test if motifs had more often a specific TF binding pattern given a specific expression cluster, than compared to the total background numbers (see Fig 7D for a diagram of the test). Results for binding patterns that did not involve a change, i.e. bound at all weeks or no weeks, were not shown in the results.

### Reduced Representation Bisulfite Sequencing

Livers from four fish per time point (three fish for week 25) were sampled and liver tissue was stored on RNAlater at −20°C. The samples were processed with Ovation RRBS Methyl-Seq System (NuGen) and bisulfite treatment was done with the Epitect Fast Bisulfite Conversion kit (Qiagen). RRBS libraries were controlled with a BioAnalyser (Agilent) machine on DNA1000 chips. Paired-end sequencing was performed by Novogene with a HiSeq X sequencing (Illumina). The mean library insert size was 168 bp and read depth was 26-40M reads.

### Alignment of bisulfite-treated reads and cytosine methylation calls

Quality trimming of reads was done with Trim Galore (v0.6.4) [81] and adapters were removed with cutadapt (v2.7) [82]. Bismark Bisulfite Mapper (v0.22.3) [81] was run with Bowtie 2 [83] against the bisulfite genome of Atlantic salmon (ICSASG_v2) [84] with the specified parameters: -q --score-min L,0,-0.2 --ignore-quals --no-mixed --no-discordant --dovetail --maxins 500. Alignment to complementary strands were ignored by default. About 40-50% of the reads were mapped to the genome. Methylated cytosines in a CpG context were extracted from the report.txt-files produced by the Bismark methylation extractor. The resulting coverage files containing methylated and unmethylated CpG loci for each sample was first filtered for known SNPs in the salmon genome then used in the analysis of differential methylation. Coverage distribution around transcription start sites (TSS) and 20 kb upstream and downstream showed that the highest coverage of reads was found nearby TSS (Fig S3B) indicating that MSPI digestion of CCGGs have resulted in enrichment around TSS, as expected by the RRBS method. Using genome annotation information, we classified CpGs according to their genomic context (Fig S3C).

### Differential methylation analysis

Samples were first organized with the R package methylKit (v1.9.4) [85] and the CpG loci were filtered by read coverage, discarding those below 10 reads per locus or more than 99.9th percentile of coverage in each sample, and those not to chromosomes. CpG loci between replicates were merged, keeping those present in at least three of the samples. The differential methylation analysis was done with an ANOVA-like analysis test of edgeR [77], contrasting the counts of methylated reads at different time points. Differentially methylated CpGs (DMC) were called with an FDR <0.05. A heatmap of the differentially methylated CpGs shows row scaled methylation percentage values of DMCs (Fig S3F).

## Data availability

Sequencing data is in the European Nucleotide database for RNA-seq (PRJEB52829), ATAC-seq (PRJEB65073), and RRBS (PRJEB60411). Code for running all steps of the analysis and generating results and figures is available on gitlab (gitlab.com/sandve-lab/GSFsmolt).

## Acknowledgments

Trond M. Kortner, from the Faculty of Veterinary Medicine, Norwegian University of Life Sciences, provided expertise during revision of the manuscript and interpretation of results.

## Supplementary material

**Fig S1.** Gene expression changes in response to salinity. Relative liver expression of genes differentially expressed (FDR <0.05) between the seawater exposed experimental group and freshwater control group at week 25. Genes are marked to the right if differentially expressed at week 25, green if higher in seawater, blue if higher in freshwater.

**Fig S2.** Principal component analysis of gene expression differences between samples. A) Similarity between all samples in the study, by principal components (PC) of gene expression. Experimental samples are colored red, and control samples for the same time points are colored gray. Labeled are potential outlier samples. B) Effect of removing sample ‘week_10_2_3’ from the PCA.

**Fig S3.** Methylated CpGs from RRBS assay. A) Similarity of samples by principal components (PC) of the read coverage of consensus CpGs. B) Genomic context of the differentially methylated CpGs. C) Heatmap of methylation values of differentially methylated CpGs during the smoltification trial. D) Density of correlation values between differentially methylated CpGs and gene expression levels, and density of correlation levels between random gene-CpG pairs.

**Table S1.** Phenotypic data of sampled Atlantic salmon. Phenotype data of Atlantic salmon used in this study. Columns provide: Unique fish identifier (*Fish ID*), week fish was sampled (*Week #*), tank number (*Tank #*), fish number (*Fish #*), date fish was sampled (*Date sampled*), fish weight (*Weight (g)*), fish length (*Length (mm)*), fish sex (*sex (M/F)*).

**Table S2.** Differentially expressed genes across smoltification. Atlantic salmon genes differentially expressed (FDR <0.05) between any time points across the smoltification experiment. Columns provide: the NCBI id for genes (*gene_id)*, available gene name (*gene_name)*, description of coded protein (*description*), and expression cluster number genes were assigned to (*deg_cluster*).

**Table S3.** Differentially expressed genes in response to photoperiod at week 10. Atlantic salmon genes differentially expressed (FDR <0.05) between week 10 experimental samples exposed prior to short photoperiod conditions, and week 10 control samples kept under continuous long photoperiod conditions. Columns provide: the NCBI id for genes (*gene_id)*, available gene name (*gene_name)*, description of coded protein (*description*), log_2_ fold change in expression of experiment versus control values (*logFC*), average expression across samples in log_2_ counts per million (logCPM), p-value of the differential expression test (*PValue*), false discovery rate adjusted p-value (*FDR*), and the time point that was tested, in this case week 10 (*week*).

**Table S4.** Differentially expressed genes in response to photoperiod at week 19. Atlantic salmon genes differentially expressed (FDR <0.05) between week 19 experimental samples exposed prior to short photoperiod conditions, and week 19 control samples kept under continuous long photoperiod conditions. Columns provide: the NCBI id for genes (*gene_id)*, available gene name (*gene_name)*, description of coded protein (*description*), log_2_ fold change in expression of experiment versus control values (*logFC*), average expression across samples in log_2_ counts per million (logCPM), p-value of the differential expression test (*PValue*), false discovery rate adjusted p-value (*FDR*), and the time point that was tested, in this case week 19 (*week*).

**Table S5.** Differentially expressed genes in response to seawater transition. Atlantic salmon genes differentially expressed (FDR <0.05) between week 25 experimental samples after transition to seawater conditions, and week 25 control samples kept under freshwater conditions. Columns provide: the NCBI ID for genes (*gene_id)*, available gene name (*gene_name)*, description of coded protein (*description*), log_2_ fold change in expression of experiment versus control values (*logFC*), average expression across samples in log_2_ counts per million (*logCPM*), p-value of the differential expression test (*PValue*), false discovery rate adjusted p-value (*FDR*).

**Table S6.** Global changes in transcription factor binding. Results from TOBIAS ATAC-seq footprinting of transcription factor binding sites (TFBS) in the Atlantic salmon genome, showing the global changes in transcription factor (TF) binding for all TF motifs tested, across the different time points of the experiment. Columns provide: output file prefix of TF name with motif ID (*output_prefix*), TF name (*name*), motif ID (*motif_id*), name of the TF’s cluster group (*cluster*), total number of TFBS (*total_tfbs*), columns for the mean score of TF binding across all TFBS for each time point (columns *week_1_mean_score* to *week_25_mean_score*), total number of bound TFBS for each time point (columns *week_10_bound* to *week_25_bound*), and the fold change followed by the p-value of the significance of the change for each pair of different time points (columns *week_1_week_10_change, week_1_week_10_pvalue*to *week_19_week_25_change, week_19_week_25_pvalue*).

**Table S7.** Changes in transcription factor binding to salinity and photoperiod. Transcription factors (TF) with significate changes in global binding of transcription factor binding sites (TFBS) between time points representing a concerted change due to experimental conditions. To photoperiod conditions; week 1 (light) vs week 10 (dark), and week 10 (dark) vs 19 (light), or to salinity conditions; week 1 (fresh) vs week 25 (sea), and week 19 (fresh) vs week 25 (sea). Columns provide: TF name (*name*), time point comparison (*comparison*), p-value of the significance of the change between time points (*pvalue*), the fold change in different of TF binding scores (*change*), the conditions compared; photoperiod or salinity (*category*), the time point where positive change means more binding (*week_A*), the time point where negative change means more binding (*week_B*), and the conditions where there is significantly more binding of the TF (*sig*).

**Table S8.** Enrichment in binding patterns of transcription factor binding sites of genes in expression clusters. Results of Fisher’s exact tests for the enrichment in different binding patterns of transcription factor binding sites (TFBS) of genes in different expression clusters. Tested for each transcription factor (TF). Columns provide: the gene expression cluster (*deg_cluster*), the binding pattern tested made up of 4 digits representing the 4 time points in chronological order with *0* equating to the TFBS not bound while *1* is bound (*binding*), the total number of TFBS with the binding pattern associated by nearest proximity to genes within the expression cluster (*count*), p-value of Fisher’s exact test (*pval*), odds ratio of test (*OR*), name of the TF motif for the TFBS (*TFBS_name*).

**Table S9.** Differentially methylated CpGs CpG sites significantly differentially methylated between time points of the experiment (FDR <0.05). Columns provide: Percentage score of the number of reads methylated at the CpG site for each time point (columns *week_1_score* to *week_25_score*), unique position of site in Atlantic salmon genome (*uniq_pos*), chromosome of site (*chr*), base position on chromosome (*locus*), log_2_ fold change in methylated read count across time points (columns *week_10_week_1_logFC* to *week_25_week_19_logFC*), average count of methylated reads across samples in log_2_ counts per million (log*CPM*), p-value for significance in change between any time points (*PValue*), false discovery rate adjusted p-value (*FDR*), associated gene’s NCBI ID (*gene_id*), position of gene transcription start site (TSS) (*tss*), gene strand position (*strand*), distance of gene TSS to CpG site (*distance*), start and end positions of gene (*gene_start, gene_end*), gene width (*gene_width*), and type of genomic feature the CpG site is located in (*genomic_feature*).

## References

1. Hoar WS. 4 The Physiology of Smolting Salmonids. In: Hoar WS, Randall DJ, editors. Fish Physiology. 11: Academic Press; 1988. p. 275-343.

2. McCormick SD, Moriyama S, Björnsson BT. Low temperature limits photoperiod control of smolting in Atlantic salmon through endocrine mechanisms. American Journal of Physiology-Regulatory, Integrative and Comparative Physiology. 2000;278(5):R1352–R61. doi: 10.1152/ajpregu.2000.278.5.R1352. PubMed PMID: 10801307.

3. Iversen M, Mulugeta T, Gellein Blikeng B, West AC, Jørgensen EH, Rød Sandven S, et al. RNA profiling identifies novel, photoperiod-history dependent markers associated with enhanced saltwater performance in juvenile Atlantic salmon. PLOS ONE. 2020;15(4):e0227496. doi: 10.1371/journal.pone.0227496.

4. McCormick SD, Regish AM, Christensen AK. Distinct freshwater and seawater isoforms of Na+/K+-ATPase in gill chloride cells of Atlantic salmon. J Exp Biol. 2009;212(24):3994–4001. doi: 10.1242/jeb.037275.

5. West AC, Mizoro Y, Wood SH, Ince LM, Iversen M, Jørgensen EH, et al. Immunologic Profiling of the Atlantic Salmon Gill by Single Nuclei Transcriptomics. Frontiers in Immunology. 2021;12. doi: 10.3389/fimmu.2021.669889.

6. Sheridan MA. Alterations in lipid metabolism accompanying smoltification and seawater adaptation of salmonid fish. Aquaculture. 1989;82(1):191–203. doi: 10.1016/0044-8486(89)90408-0.

7. Sheridan MA, Woo NYS, Bern HA. Changes in the rates of glycogenesis, glycogenolysis, lipogenesis, and lipolysis in selected tissues of the coho salmon (Oncorhynchus kisutch) associated with parr-smolt transformation. Journal of Experimental Zoology. 1985;236(1):35–44. doi: 10.1002/jez.1402360106.

8. McCormick SD. 5 - Smolt Physiology and Endocrinology. In: McCormick SD, Farrell AP, Brauner CJ, editors. Fish Physiology. 32: Academic Press; 2012. p. 199-251.

9. Solbakken VA, Hansen T, Stefansson SO. Effects of photoperiod and temperature on growth and parr-smolt transformation in Atlantic salmon (Salmo salar L.) and subsequent performance in seawater. Aquaculture. 1994;121(1):13–27. doi: 10.1016/0044-8486(94)90004-3.

10. Strand JET, Hazlerigg D, Jørgensen EH. Photoperiod revisited: is there a critical day length for triggering a complete parr–smolt transformation in Atlantic salmon Salmo salar? J Fish Biol. 2018;93(3):440–8. doi: 10.1111/jfb.13760.

11. Iversen M, Mulugeta T, West AC, Jørgensen EH, Martin SAM, Sandve SR, et al. Photoperiod-dependent developmental reprogramming of the transcriptional response to seawater entry in Atlantic salmon (Salmo salar). G3 Genes|Genomes|Genetics. 2021;11(4). doi: 10.1093/g3journal/jkab072.

12. Bell JG, Tocher DR, Farndale BM, Cox DI, McKinney RW, Sargent JR. The effect of dietary lipid on polyunsaturated fatty acid metabolism in Atlantic salmon (Salmo salar) undergoing parr-smolt transformation. Lipids. 1997;32(5):515–25. doi: 10.1007/s11745-997-0066-4.

13. Tocher DR, Bell JG, Dick JR, Henderson RJ, McGhee F, Michell D, et al. Polyunsaturated fatty acid metabolism in Atlantic salmon (Salmo salar) undergoing parr-smolt transformation and the effects of dietary linseed and rapeseed oils. Fish Physiol Biochem. 2000;23(1):59–73. doi: 10.1023/A:1007807201093.

14. Gillard G, Harvey TN, Gjuvsland A, Jin Y, Thomassen M, Lien S, et al. Life-stage-associated remodelling of lipid metabolism regulation in Atlantic salmon. Mol Ecol. 2018;27(5):1200–13. doi: 10.1111/mec.14533.

15. Li LO, Klett EL, Coleman RA. Acyl-CoA synthesis, lipid metabolism and lipotoxicity. Biochim Biophys Acta. 2010;1801(3):246–51. doi: 10.1016/j.bbalip.2009.09.024.

16. Yen C-LE, Stone SJ, Koliwad S, Harris C, Farese RV. Thematic Review Series: Glycerolipids. DGAT enzymes and triacylglycerol biosynthesis. J Lipid Res. 2008;49(11):2283–301. doi: 10.1194/jlr.R800018-JLR200.

17. Sul HS, Smith S. CHAPTER 6 - Fatty acid synthesis in eukaryotes. In: Vance DE, Vance JE, editors. Biochemistry of Lipids, Lipoproteins and Membranes (Fifth Edition). San Diego: Elsevier; 2008. p. 155–90.

18. Grevengoed TJ, Klett EL, Coleman RA. Acyl-CoA Metabolism and Partitioning. Annu Rev Nutr. 2014;34(1):1–30. doi: 10.1146/annurev-nutr-071813-105541. PubMed PMID: 24819326.

19. Tillander V, Alexson SEH, Cohen DE. Deactivating Fatty Acids: Acyl-CoA Thioesterase-Mediated Control of Lipid Metabolism. Trends in Endocrinology & Metabolism. 2017;28(7):473–84. doi: 10.1016/j.tem.2017.03.001.

20. Jones G, Strugnell SA, DeLuca HF. Current Understanding of the Molecular Actions of Vitamin D. Physiol Rev. 1998;78(4):1193–231. doi: 10.1152/physrev.1998.78.4.1193. PubMed PMID: 9790574.

21. Pierens SL, Fraser DR. The origin and metabolism of vitamin D in rainbow trout. The Journal of steroid biochemistry and molecular biology. 2015;145:58–64. doi: 10.1016/j.jsbmb.2014.10.005. PubMed PMID: 25305412.

22. Meinsohn M-C, Smith OE, Bertolin K, Murphy BD. The Orphan Nuclear Receptors Steroidogenic Factor-1 and Liver Receptor Homolog-1: Structure, Regulation, and Essential Roles in Mammalian Reproduction. Physiol Rev. 2019;99(2):1249–79. doi: 10.1152/physrev.00019.2018. PubMed PMID: 30810078.

23. Olbrot M, Rud J, Moss LG, Sharma A. Identification of β-cell-specific insulin gene transcription factor RIPE3b1 as mammalian MafA. Proceedings of the National Academy of Sciences. 2002;99(10):6737–42. doi: doi:10.1073/pnas.102168499.

24. Yerra VG, Drosatos K. Specificity Proteins (SP) and Krppel-like Factors (KLF) in Liver Physiology and Pathology. International Journal of Molecular Sciences. 2023;24(5):4682. PubMed PMID: doi:10.3390/ijms24054682.

25. Kim YH, Lazar MA. Transcriptional Control of Circadian Rhythms and Metabolism: A Matter of Time and Space. Endocr Rev. 2020;41(5):707–32. doi: 10.1210/endrev/bnaa014.

26. Zhu B. Decoding the function and regulation of the mammalian 12-h clock. Journal of Molecular Cell Biology. 2020;12(10):752–8. doi: 10.1093/jmcb/mjaa021.

27. Lee C-H, Olson P, Hevener A, Mehl I, Chong L-W, Olefsky JM, et al. PPARδ regulates glucose metabolism and insulin sensitivity. Proceedings of the National Academy of Sciences. 2006;103(9):3444–9. doi: doi:10.1073/pnas.0511253103.

28. Carmona-Antoñanzas G, Tocher DR, Martinez-Rubio L, Leaver MJ. Conservation of lipid metabolic gene transcriptional regulatory networks in fish and mammals. Gene. 2014;534(1):1–9. doi: 10.1016/j.gene.2013.10.040.

29. Huang K-W, Reebye V, Czysz K, Ciriello S, Dorman S, Reccia I, et al. Liver Activation of Hepatocellular Nuclear Factor-4α by Small Activating RNA Rescues Dyslipidemia and Improves Metabolic Profile. Molecular Therapy - Nucleic Acids. 2020;19:361–70. doi: 10.1016/j.omtn.2019.10.044.

30. Mullur R Fau - Liu Y-Y, Liu Yy Fau - Brent GA, Brent GA. Thyroid hormone regulation of metabolism. (1522-1210 (Electronic)).

31. Walker M, Li Y, Morales-Hernandez A, Qi Q, Parupalli C, Brown SA, et al. An NFIX-mediated regulatory network governs the balance of hematopoietic stem and progenitor cells during hematopoiesis. Blood Advances. 2022. doi: 10.1182/bloodadvances.2022007811.

32. Hernandez JM, Floyd DH, Weilbaecher KN, Green PL, Boris-Lawrie K. Multiple facets of junD gene expression are atypical among AP-1 family members. Oncogene. 2008;27(35):4757–67. doi: 10.1038/onc.2008.120.

33. Hess J, Angel P, Schorpp-Kistner M. AP-1 subunits: quarrel and harmony among siblings. J Cell Sci. 2004;117(25):5965–73. doi: 10.1242/jcs.01589.

34. Nisembaum LG, Martin P, Lecomte F, Falcón J. Melatonin and osmoregulation in fish: A focus on Atlantic salmon Salmo salar smoltification. J Neuroendocrinol. 2021;33(3):e12955. doi: 10.1111/jne.12955.

35. Frey-Jakobs S, Hartberger JM, Fliegauf M, Bossen C, Wehmeyer ML, Neubauer JC, et al. ZNF341 controls STAT3 expression and thereby immunocompetence. Science Immunology. 2018;3(24):eaat4941. doi: doi:10.1126/sciimmunol.aat4941.

36. Golson ML, Kaestner KH. Fox transcription factors: from development to disease. Development. 2016;143(24):4558–70. doi: 10.1242/dev.112672.

37. Hannenhalli S, Kaestner KH. The evolution of Fox genes and their role in development and disease. Nature Reviews Genetics. 2009;10(4):233–40. doi: 10.1038/nrg2523.

38. Weiss RJ, Spahn PN, Toledo AG, Chiang AWT, Kellman BP, Li J, et al. ZNF263 is a transcriptional regulator of heparin and heparan sulfate biosynthesis. Proceedings of the National Academy of Sciences. 2020;117(17):9311–7. doi: doi:10.1073/pnas.1920880117.

39. Yoon HS, Scharer CD, Majumder P, Davis CW, Butler R, Zinzow-Kramer W, et al. ZBTB32 Is an Early Repressor of the CIITA and MHC Class II Gene Expression during B Cell Differentiation to Plasma Cells. The Journal of Immunology. 2012;189(5):2393–403. doi: 10.4049/jimmunol.1103371.

40. Arroyo JD, Jourdain AA, Calvo SE, Ballarano CA, Doench JG, Root DE, et al. A Genome-wide CRISPR Death Screen Identifies Genes Essential for Oxidative Phosphorylation. Cell Metabolism. 2016;24(6):875–85. doi: 10.1016/j.cmet.2016.08.017.

41. Su J, Morgani SM, David CJ, Wang Q, Er EE, Huang Y-H, et al. TGF-β orchestrates fibrogenic and developmental EMTs via the RAS effector RREB1. Nature. 2020;577(7791):566-71. doi: 10.1038/s41586-019-1897-5.

42. Thiagalingam A, Bustros Ad, Borges M, Jasti R, Compton D, Diamond L, et al. RREB-1, a Novel Zinc Finger Protein, Is Involved in the Differentiation Response to Ras in Human Medullary Thyroid Carcinomas. Mol Cell Biol. 1996;16(10):5335–45. doi: 10.1128/MCB.16.10.5335.

43. Wung BS, Cheng JJ, Chao YJ, Hsieh HJ, Wang DL. Modulation of Ras/Raf/Extracellular Signal–Regulated Kinase Pathway by Reactive Oxygen Species Is Involved in Cyclic Strain–Induced Early Growth Response-1 Gene Expression in Endothelial Cells. Circul Res. 1999;84(7):804–12. doi: doi:10.1161/01.RES.84.7.804.

44. Reed BD, Charos AE, Szekely AM, Weissman SM, Snyder M. Genome-Wide Occupancy of SREBP1 and Its Partners NFY and SP1 Reveals Novel Functional Roles and Combinatorial Regulation of Distinct Classes of Genes. PLoS Genet. 2008;4(7):e1000133. doi: 10.1371/journal.pgen.1000133.

45. Versteyhe S, Klaproth B, Borup R, Palsgaard J, Jensen M, Gray S, et al. IGF-I, IGF-II, and Insulin Stimulate Different Gene Expression Responses through Binding to the IGF-I Receptor. Frontiers in Endocrinology. 2013;4. doi: 10.3389/fendo.2013.00098.

46. Ye D, Lou G, Zhang T, Dong F, Liu Y. MiR-17 family-mediated regulation of Pknox1 influences hepatic steatosis and insulin signaling. J Cell Mol Med. 2018;22(12):6167–75. doi: 10.1111/jcmm.13902.

47. Hunter AL, Pelekanou CE, Adamson A, Downton P, Barron NJ, Cornfield T, et al. Nuclear receptor REVERBα is a state-dependent regulator of liver energy metabolism. Proceedings of the National Academy of Sciences. 2020;117(41):25869–79. doi: doi:10.1073/pnas.2005330117.

48. Oishi Y, Manabe I. Krüppel-Like Factors in Metabolic Homeostasis and Cardiometabolic Disease. Frontiers in Cardiovascular Medicine. 2018;5. doi: 10.3389/fcvm.2018.00069.

49. Pisam M, Prunet P, Boeuf G, Jrambourg A. Ultrastructural features of chloride cells in the gill epithelium of the atlantic salmon, Salmo salar, and their modifications during smoltification. American Journal of Anatomy. 1988;183(3):235–44. doi: 10.1002/aja.1001830306.

50. Evans DH, Piermarini PM, Choe KP. The Multifunctional Fish Gill: Dominant Site of Gas Exchange, Osmoregulation, Acid-Base Regulation, and Excretion of Nitrogenous Waste. Physiol Rev. 2005;85(1):97–177. doi: 10.1152/physrev.00050.2003. PubMed PMID: 15618479.

51. Kiilerich P, Kristiansen K, Madsen SS. Cortisol regulation of ion transporter mRNA in Atlantic salmon gill and the effect of salinity on the signaling pathway. J Endocrinol. 2007;194(2):417–27. doi: 10.1677/JOE-07-0185.

52. Nilsen TO, Ebbesson LOE, Madsen SS, McCormick SD, Andersson E, Björnsson BrT, et al. Differential expression of gill Na+,K+-ATPaseα- and β-subunits, Na+,K+,2Cl-cotransporter and CFTR anion channel in juvenile anadromous and landlocked Atlantic salmon Salmo salar. J Exp Biol. 2007;210(16):2885–96. doi: 10.1242/jeb.002873.

53. Tipsmark CK, Jørgensen C, Brande-Lavridsen N, Engelund M, Olesen JH, Madsen SS. Effects of cortisol, growth hormone and prolactin on gill claudin expression in Atlantic salmon. Gen Comp Endocrinol. 2009;163(3):270–7. doi: 10.1016/j.ygcen.2009.04.020.

54. Itokazu Y, Käkelä R, Piironen J, Guan XL, Kiiskinen P, Vornanen M. Gill tissue lipids of salmon (Salmo salar L.) presmolts and smolts from anadromous and landlocked populations. Comparative Biochemistry and Physiology Part A: Molecular & Integrative Physiology. 2014;172:39–45. doi: 10.1016/j.cbpa.2014.01.020.

55. Ackman RG. Characteristics of the fatty acid composition and biochemistry of some fresh-water fish oils and lipids in comparison with marine oils and lipids. Comparative Biochemistry and Physiology. 1967;22(3):907–22. doi: 10.1016/0010-406X(67)90781-5.

56. Sheridan MA, Allen WV, Kerstetter TH. Changes in the fatty acid composition of steelhead trout, Salmo gairdnerii Richardson, associated with parr-smolt transformation. Comparative Biochemistry and Physiology Part B: Comparative Biochemistry. 1985;80(4):671–6. doi: 10.1016/0305-0491(85)90444-4.

57. Björnsson BT, Thorarensen H, Hirano T, Ogasawara T, Kristinsson JB. Photoperiod and temperature affect plasma growth hormone levels, growth, condition factor and hypoosmoregulatory ability of juvenile Atlantic salmon (Salmo salar) during parr-smolt transformation. Aquaculture. 1989;82(1):77–91. doi: 10.1016/0044-8486(89)90397-9.

58. Berge ÅI, Berg A, Barnung T, Hansen T, Fyhn HJ, Stefansson SO. Development of salinity tolerance in underyearling smolts of Atlantic salmon (Salmo salar) reared under different photoperiods. Can J Fish Aquat Sci. 1995;52(2):243–51. doi: 10.1139/f95-024.

59. Duston J, Saunders RL. Advancing smolting to autumn in age 0+ Atlantic salmon by photoperiod, and long-term performance in sea water. Aquaculture. 1995;135(4):295–309. doi: 10.1016/0044-8486(95)01034-3.

60. McCormick SD, Björnsson BT, Sheridan M, Eilerlson C, Carey JB, O’Dea M. Increased daylength stimulates plasma growth hormone and gill Na+, K+-ATPase in Atlantic salmon (Salmo salar). Journal of Comparative Physiology B. 1995;165(4):245–54. doi: 10.1007/BF00367308.

61. McCormick SD, Shrimpton JM, Moriyama S, Björnsson BT. Differential hormonal responses of Atlantic salmon parr and smolt to increased daylength: A possible developmental basis for smolting. Aquaculture. 2007;273(2):337–44. doi: 10.1016/j.aquaculture.2007.10.015.

62. Duncan NJ, Bromage N. The effect of different periods of constant short days on smoltification in juvenile Atlantic salmon (Salmo salar). Aquaculture. 1998;168(1):369–86. doi: 10.1016/S0044-8486(98)00363-9.

63. Handeland SO, Stefansson SO. Photoperiod control and influence of body size on off-season parr–smolt transformation and post-smolt growth. Aquaculture. 2001;192(2):291–307. doi: 10.1016/S0044-8486(00)00457-9.

64. Stefansson SO, Nilsen TO, Ebbesson LOE, Wargelius A, Madsen SS, Björnsson BT, et al. Molecular mechanisms of continuous light inhibition of Atlantic salmon parr–smolt transformation. Aquaculture. 2007;273(2):235–45. doi: 10.1016/j.aquaculture.2007.10.005.

65. Ross AW, Webster CA, Mercer JG, Moar KM, Ebling FJ, Schuhler S, et al. Photoperiodic Regulation of Hypothalamic Retinoid Signaling: Association of Retinoid X Receptor γ with Body Weight. Endocrinology. 2004;145(1):13–20. doi: 10.1210/en.2003-0838.

66. Verma R, Haldar C. Photoperiodic modulation of thyroid hormone receptor (TR-α), deiodinase-2 (Dio-2) and glucose transporters (GLUT 1 and GLUT 4) expression in testis of adult golden hamster, Mesocricetus auratus. J Photochem Photobiol B: Biol. 2016;165:351–8. doi: 10.1016/j.jphotobiol.2016.10.036.

67. Morán P, Pérez-Figueroa A. Methylation changes associated with early maturation stages in the Atlantic salmon. BMC Genet. 2011;12(1):86. doi: 10.1186/1471-2156-12-86.

68. Mohamed AR, Naval-Sanchez M, Menzies M, Evans B, King H, Reverter A, et al. Leveraging transcriptome and epigenome landscapes to infer regulatory networks during the onset of sexual maturation. BMC Genomics. 2022;23(1):413. doi: 10.1186/s12864-022-08514-8.

69. Riviere G, He Y, Tecchio S, Crowell E, Gras M, Sourdaine P, et al. Dynamics of DNA methylomes underlie oyster development. PLoS Genet. 2017;13(6):e1006807. doi: 10.1371/journal.pgen.1006807.

70. Gegner J, Vogel H, Billion A, Förster F, Vilcinskas A. Complete Metamorphosis in Manduca sexta Involves Specific Changes in DNA Methylation Patterns. Frontiers in Ecology and Evolution. 2021;9. doi: 10.3389/fevo.2021.646281.

71. Cardoso-Júnior CAM, Yagound B, Ronai I, Remnant EJ, Hartfelder K, Oldroyd BP. DNA methylation is not a driver of gene expression reprogramming in young honey bee workers. Mol Ecol. 2021;30(19):4804–18. doi: 10.1111/mec.16098.

72. Hayashi AA, Proud CG. The rapid activation of protein synthesis by growth hormone requires signaling through mTOR. American Journal of Physiology-Endocrinology and Metabolism. 2007;292(6):E1647–E55. doi: 10.1152/ajpendo.00674.2006. PubMed PMID: 17284572.

73. Laplante M, Sabatini DM. Regulation of mTORC1 and its impact on gene expression at a glance. (1477-9137 (Electronic)).

74. Handeland SO, Björnsson Arnesen AM, Stefansson SO. Seawater adaptation and growth of post-smolt Atlantic salmon (Salmo salar) of wild and farmed strains. Aquaculture. 2003;220(1):367–84. doi: 10.1016/S0044-8486(02)00508-2.

75. Sakamoto T, McCormick SD, Hirano T. Osmoregulatory actions of growth hormone and its mode of action in salmonids: A review. Fish Physiol Biochem. 1993;11(1):155–64. doi: 10.1007/BF00004562.

76. Usher ML, Talbot C, Eddy FB. Effects of transfer to seawater on growth and feeding in Atlantic salmon smolts (Salmo salar L.). Aquaculture. 1991;94(4):309–26. doi: 10.1016/0044-8486(91)90176-8.

77. Robinson MD, McCarthy DJ, Smyth GK. edgeR: a Bioconductor package for differential expression analysis of digital gene expression data. Bioinformatics. 2009;26(1):139–40. doi: 10.1093/bioinformatics/btp616.

78. Buenrostro JD, Giresi PG, Zaba LC, Chang HY, Greenleaf WJ. Transposition of native chromatin for fast and sensitive epigenomic profiling of open chromatin, DNA-binding proteins and nucleosome position. Nat Methods. 2013;10(12):1213–8. doi: 10.1038/nmeth.2688.

79. Buenrostro JD, Wu B, Chang HY, Greenleaf WJ. ATAC-seq: A Method for Assaying Chromatin Accessibility Genome-Wide. Current Protocols in Molecular Biology. 2015;109(1):21.9.1-.9.9. doi: 10.1002/0471142727.mb2129s109.

80. Bentsen M, Goymann P, Schultheis H, Klee K, Petrova A, Wiegandt R, et al. ATAC-seq footprinting unravels kinetics of transcription factor binding during zygotic genome activation. Nature Communications. 2020;11(1):4267. doi: 10.1038/s41467-020-18035-1.

81. Krueger F, Andrews SR. Bismark: a flexible aligner and methylation caller for Bisulfite-Seq applications. Bioinformatics. 2011;27(11):1571–2. doi: 10.1093/bioinformatics/btr167.

82. Martin M. Cutadapt removes adapter sequences from high-throughput sequencing reads. EMBnetjournal; Vol 17, No 1: Next Generation Sequencing Data AnalysisDO - 1014806/ej171200. 2011.

83. Langmead B, Salzberg SL. Fast gapped-read alignment with Bowtie 2. Nat Methods. 2012;9(4):357–9. doi: 10.1038/nmeth.1923.

84. Lien S, Koop BF, Sandve SR, Miller JR, Kent MP, Nome T, et al. The Atlantic salmon genome provides insights into rediploidization. Nature. 2016;533(7602):200-5. doi: 10.1038/nature17164.

85. Akalin A, Kormaksson M, Li S, Garrett-Bakelman FE, Figueroa ME, Melnick A, et al. methylKit: a comprehensive R package for the analysis of genome-wide DNA methylation profiles. Genome Biology. 2012;13(10):R87. doi: 10.1186/gb-2012-13-10-r87.

